# DeepMM: Identify and correct Metagenome Misassemblies with deep learning

**DOI:** 10.1101/2025.02.07.637187

**Authors:** Yi Ding, Jin Xiao, Bohao Zou, Chao Yang, Lu Zhang

## Abstract

Accurate metagenomic assemblies are essential for constructing reliable metagenome-assembled genomes (MAGs). However, the complexity of microbial genomes continues to pose challenges for accurate assembly. Current reference-free assembly evaluation tools primarily rely on hand-crafted features and suffer from poor generalization across different metagenomic data. To address these limitations, we propose DeepMM, a novel deep learning-based visual model designed for the identification and correction of metagenomic misassemblies. DeepMM transforms alignments between assemblies and reads into a multi-channel image for misassembly feature learning and applies contrastive learning to bring different views of misassemblies closer. Furthermore, DeepMM offers a fine-tuning process to match different sequencer data. Our results show that DeepMM outperforms state-of-the-art methods in identifying misassemblies, achieving the highest AUPRC score in five CAMI datasets. DeepMM provides accurate correction of misassemblies, significantly improving downstream binning results, increasing the number of near-complete MAGs from 905 to 1006 in a large real metagenomic sequencing dataset derived from a diarrhea-predominant Irritable Bowel Syndrome (IBS-D) cohort.

## 1 Introduction

Next-generation metagenome sequencing offers an accessible technology for metagenome-assembled genomes(MAGs) construction, which plays an important role in unlocking the dark matter secrets of microbial communities [1, 2]. Despite advances in current assembly algorithms, the intrinsic complexity of microbial genomes, such as inter- and intra-genomic repeat regions[3], continues to pose challenges, often leading to misassembly. Misassembly refers to the erroneous reconstruction of genome sequences during the assembly process, frequently resulting in chimeric sequences that combine fragments from distinct microbial genomes. These inaccuracies can substantially undermine the robustness and validity of downstream ecological and evolutionary analyses. Consequently, accurate metagenomic assemblies are imperative for the robust downstream analysis of MAGs.

Owing to the scarcity of reference data within metagenomic databases and leveraging the reduced expenses associated with NGS techniques[4], recent reference-free strategies have advanced in evaluating metagenomic assemblies. These strategies model misassembly by calculating descriptive statistics of alignment signals derived from the alignment of short reads against assemblies. Alignment signals that are commonly used to model misassembly include read depth (the number of reads that map to a genome region), discordant read pairs (read pairs from the same fragment whose mapping deviates in distance or orientation), and clipped reads (reads that have partial alignments to the assembly)[5]. These signals are then combined into a sophisticated statistical or machine learning model to learn and detect misassembly. ALE[6], a statistical model, utilizes Bayesian probability to describe the likelihood of each position to measure the quality of assemblies; metaMIC[5], a machine learning-based tool, harnesses a range of descriptive statistics of alignment features from alignments to detect and correct misassemblies. Deep learning, with its capacity to autonomously learn complex representations from extensive labeled datasets without expert guidance, presents a compelling approach for broadly identifying misassemblies. DeepMAsED [7] and ResMiCo [8], two deep learning-based tools, leverage features analogous to those used by metaMIC, which are systematically transformed into structured formats to facilitate misassembly detection. However, these methodologies show a substantial dependence on features engineered by manual curation, which inherently limits the scope of detectable patterns within metagenomic data. These methods demonstrate limited generalizability, potentially due to their reliance on single-view feature representations of misassemblies and their inability to accommodate varying sequencing data with different sequencer parameters.

Here, we introduce DeepMM, a deep learning-based visual model for identifying and correcting metagenomic misassemblies. DeepMM employs a refined alignment purification process to ensure robust feature extraction. By reformulating misassembly detection as an image classification task, DeepMM interprets alignments as multi-channel images, where misassemblies represent distinct visual patterns. This approach, assisted with contrastive learning to match different views of misassembly representations, enables DeepMM to effectively identify misassemblies. Furthermore, a fine-tuning process facilitates adaptation to various sequencing data characteristics. Evaluation of simulated and real metagenomic datasets demonstrates that DeepMM surpasses existing methods in misassembly identification and correction accuracy, leading to improved downstream analyses such as binning.

## 2 Results

### 2.1 Overflow of DeepMM

At a high level, assemblies are divided into sliding windows of length *l*, excluding the outermost 10% of basepairs (capped at 1kbps) at each end. For each postion in a sliding window, six features are extracted to help distinguish correctly assembled sequences from chimeric contig breakpoints: 1. Clipped reads, the total number of reads clipped at position *i*; 2. Inverted orientation reads, the total number of read pairs covering postion *i* that have the same orientation; 3. Translocated read pairs, the total number of reads covering position *i* whose mate pairs are aligned to a different contig; 4. Insert size, the average insert size of reads pairs covering position *i*; 5. Read depth, the total number of reads aligned to position *i*; 6. Read depth difference, the difference in read depth between position *i* and *j* in the same window. For each window, the features for all positions are collected and transferred into a symmetric feature map of length *l*. The first five features result in non-zero diagonal elements, as they are calculated for individual positions, where the last feature has non-zero off-diagnal elements, reflecting the depth differences between positions (Fig.1). These feature maps are stacked into a 6-channel image, where pixel intensities are converted from values in feature maps. The images are passed through the model to predict whether the window contains breakpoints of assemblies based on image embeddings. During training, we employ data augmentation by constructing four types of windows from each original window for contrastive learning, thereby maximizing the similarity of different views of a misassembly. Specifically, for a given window, we generate left-shifted and right-shifted views based on the midpoint of the original window, as well as a reversed view and a sub-sampled view by sub-sampling the aligned reads. After learning the similarity of different views and feature representations of a misassembly, the model can predict whether a window contains a misassembly. Misassembly correction is then performed based on these predictions (Fig.2). Additionally, DeepMM can be fine-tuned with different sequencing data to achieve better performance (see ‘Methods’).

**Figure 1.**
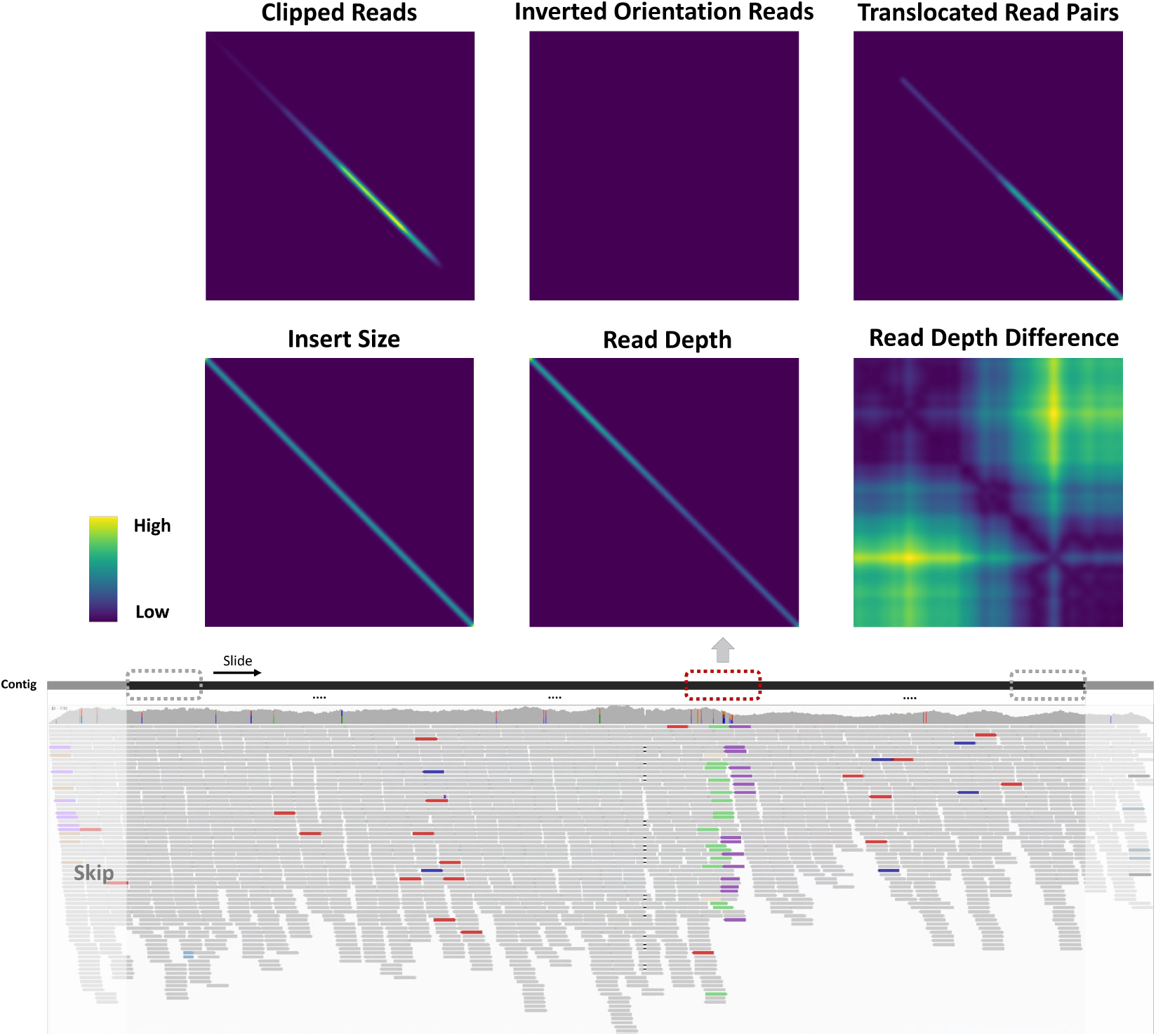
Construction of multi-channel image feature representation

**Figure 2.**
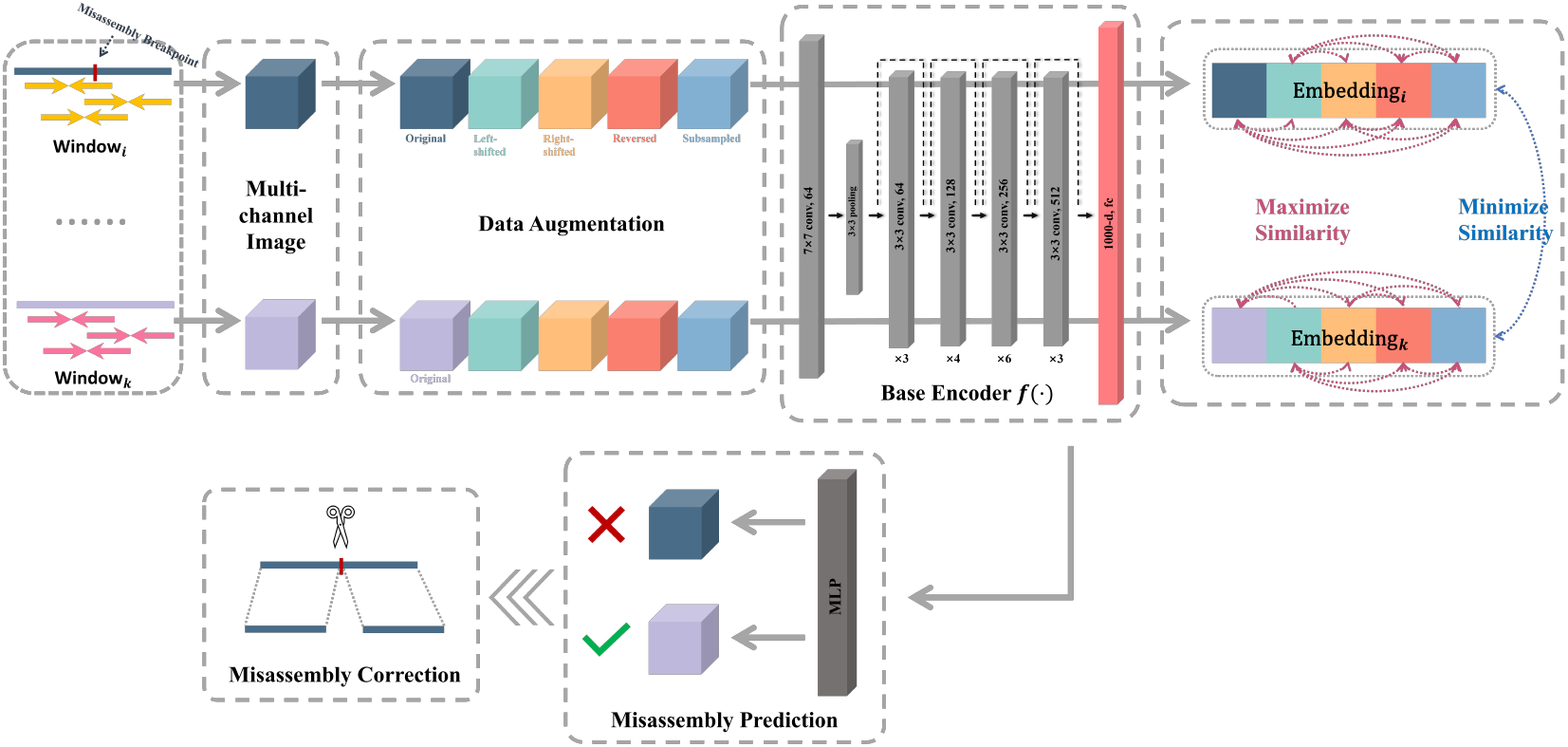
Model structure of DeepMM

### 2.2 Identifying misassemblies in simulated metagenomic datasets

To evaluate the performance in identifying misassemblies, we tested DeepMM on simulated metagenomic datasets from CAMI (the Critical Assessment of Metagenome Interpretation) [9], which comprise various microbes. We used MEGAHIT [10] to assemble reads from CAMI datasets and obtained the ground truth misassemblies through MetaQUAST [11]. We first challenged DeepMM with Medium- and High-complexity CAMI datasets to assess how dataset complexity affects DeepMM’s accuracy. Compared with the CAMI1-Medium dataset, DeepMM performed better on the CAMI1-High dataset, demonstrating its effectiveness for high microbial diversity communities and significantly outperforming existing tools by achieving the highest AUPRC, as shown in Fig. 3a,b. We also evaluated DeepMM on CAMI2-Gut, CAMI2-Oral, and CAMI2-Skin to assess its performance across different human body sites. As illustrated in Fig. 3c, d, and e, DeepMM maintains high precision across the recall threshold, delivering a consistent and reliable balance between precision and recall, whereas other tools show steep declines or instability at higher recall rates.

**Figure 3.**
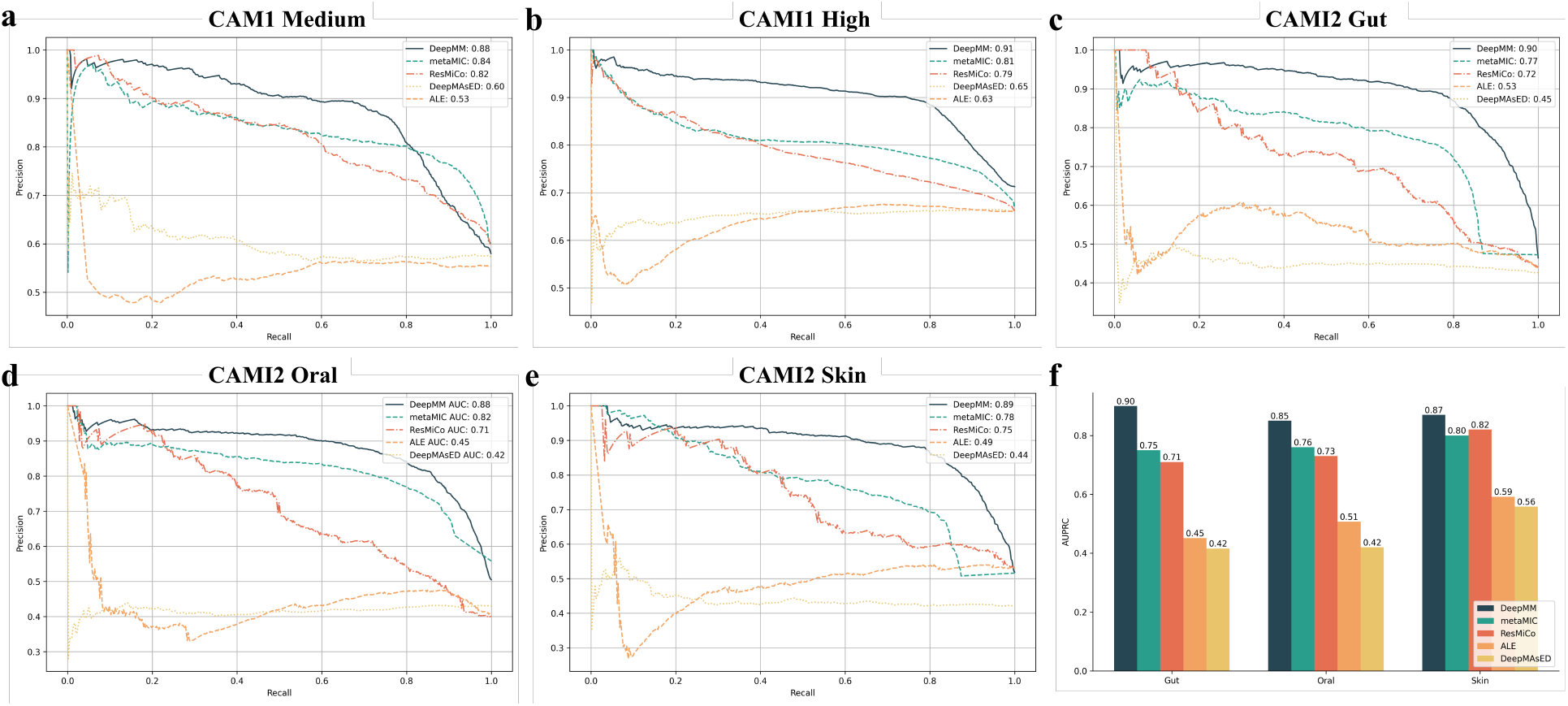
AUPRC scores of identifying misassemblies in five CAMI datasets

Given that DeepMM is trained on datasets where assemblies were constructed by MEGAHIT, we further investigated its generalization across different assemblers directly. We assembled reads of CAMI2-Gut, CAMI2-Oral, and CAMI2-Skin with metaSPAdes [12]. As shown in Fig. 3f, DeepMM consistently achieves the highest AUPRC scores across all three datasets, outperforming metaMIC, ResMiCo, ALE, and DeepMasED. The consistent advantage of DeepMM across varied assemblers underscores its reliability and robustness in handling metagenomic data assembled using different tools.

### 2.3 Accurate correcting misassemblies improves binning results

Since only metaMIC provides misassembly correction, we only used metaMIC with default settings for the following experiments. We first calculated the F1-score in five CAMI datasets. As shown in Fig. 4, DeepMM achieves higher F1 scores in all datasets, showing the superiority of DeepMM in identifying misassemblies. After identifying misassemblies, DeepMM can provide a correction process by splitting the misassemblies into shorter sequences at the breakpoints. To see how the correction of splitting misassemblies at breakpoints by DeepMM will influence downstream analyses, we binned the assemblies in all CAMI datasets using SemiBin2[13] and estimated by CheckM2[14]. As shown in Fig. 5, we collected the reconstructed MAGs with different completeness (contamination *<* 5%), and the results illustrate the impact of misassembly correction on binning completeness. DeepMM achieves superior results on near-complete bins (completeness *>*90%) compared to metaMIC with higher F1-score in misassembly identification. For example, in the High-complexity dataset, DeepMM helped recover 133 bins with *>*90% completeness, surpassing metaMIC. These results demonstrate that correcting misassemblies by DeepMM significantly enhances the quality of downstream binning, particularly in complex microbial communities.

**Figure 4.**
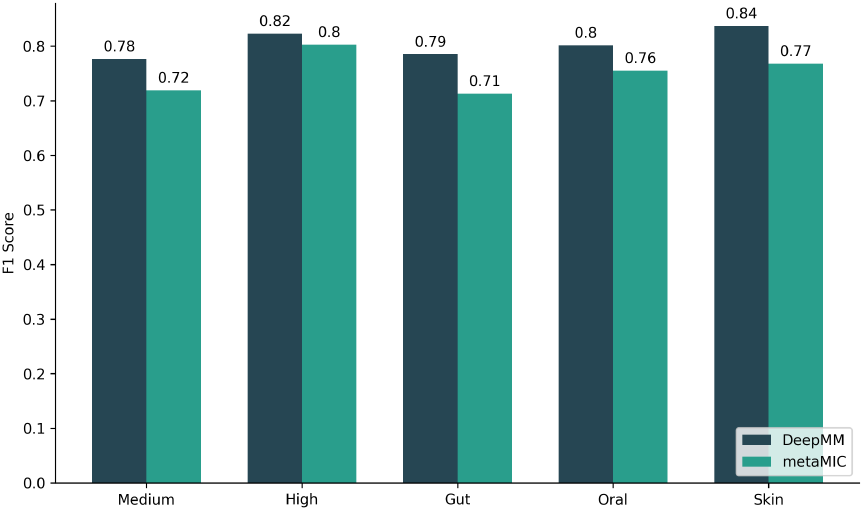
F1 score of misassembly identification in five CAMI datasets

**Figure 5.**
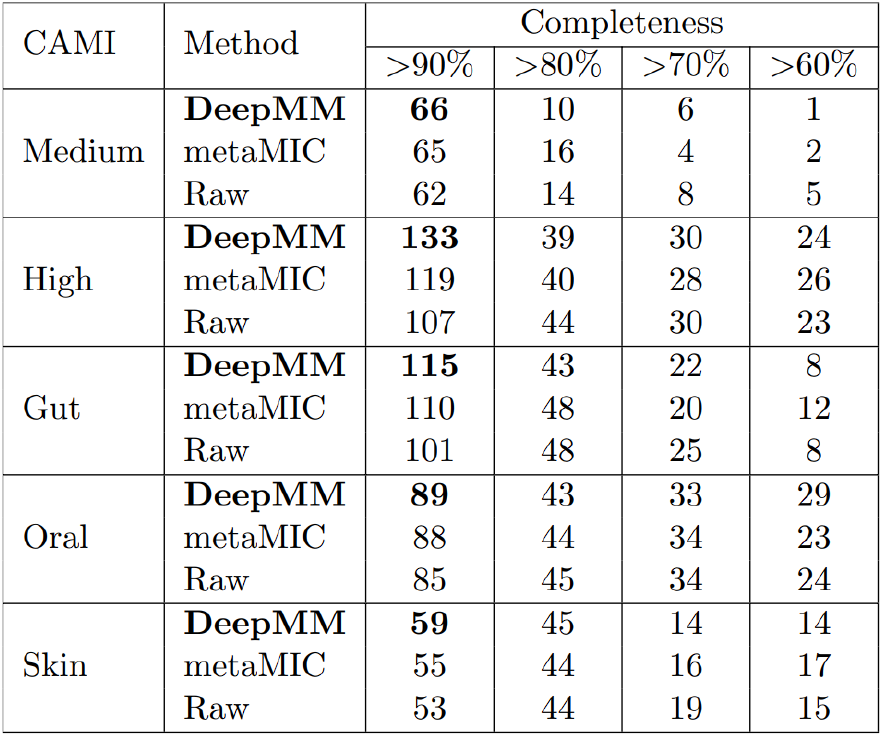
Increased the number of near-complete reconstructed bins in five CAMI datasets

### 2.4 Applying DeepMM to real metagenomic data

We applied DeepMM to a large real metagenomic sequencing dataset derived from a diarrhea-predominant Irritable Bowel Syndrome (IBS-D) cohort, which includes 290 patients and 89 healthy controls [15]. We randomly selected approximately fifty samples from the dataset to evaluate DeepMM. There are 359,610 assemblies in the randomly selected samples, and DeepMM corrected 63,043 assemblies, accounting for 17.5%. Our analysis revealed that DeepMM significantly enhanced the reconstruction of near-complete microbial genomes from IBS-D metagenomic data. Specifically, DeepMM correction facilitated the reconstruction of 1,006 bins with over 90% completeness, surpassing metaMIC, which reconstructed 950 bins. These results demonstrate DeepMM’s superior misassembly identification ability, aiding in the reconstruction of high-quality microbial genomes from complex metagenomic datasets.

As reference genome sequences are absent in real data, we used a real metagenomic sequencing dataset from gut microbiomes [16], which includes Next-generation Sequencing (NGS) data and Oxford Nanopore Technology (ONT) sequencing data, for misassembly prediction validation. As shown in Fig. 6, we aligned the NGS reads and ONT reads back to the NGS assembly. A sharp peak in the misassembly score at approximately the 13 kb region indicates a potential misassembly error, which is further validated by ONT read alignments. In this region, ONT reads display numerous disruptions, including clipped reads, mismatched bases, and indels, confirming the predicted misassembly. This case demonstrates the effectiveness of DeepMM in real data, while ONT validation provides robust evidence to resolve and verify structural inconsistencies in the assembly.

**Figure 6.**
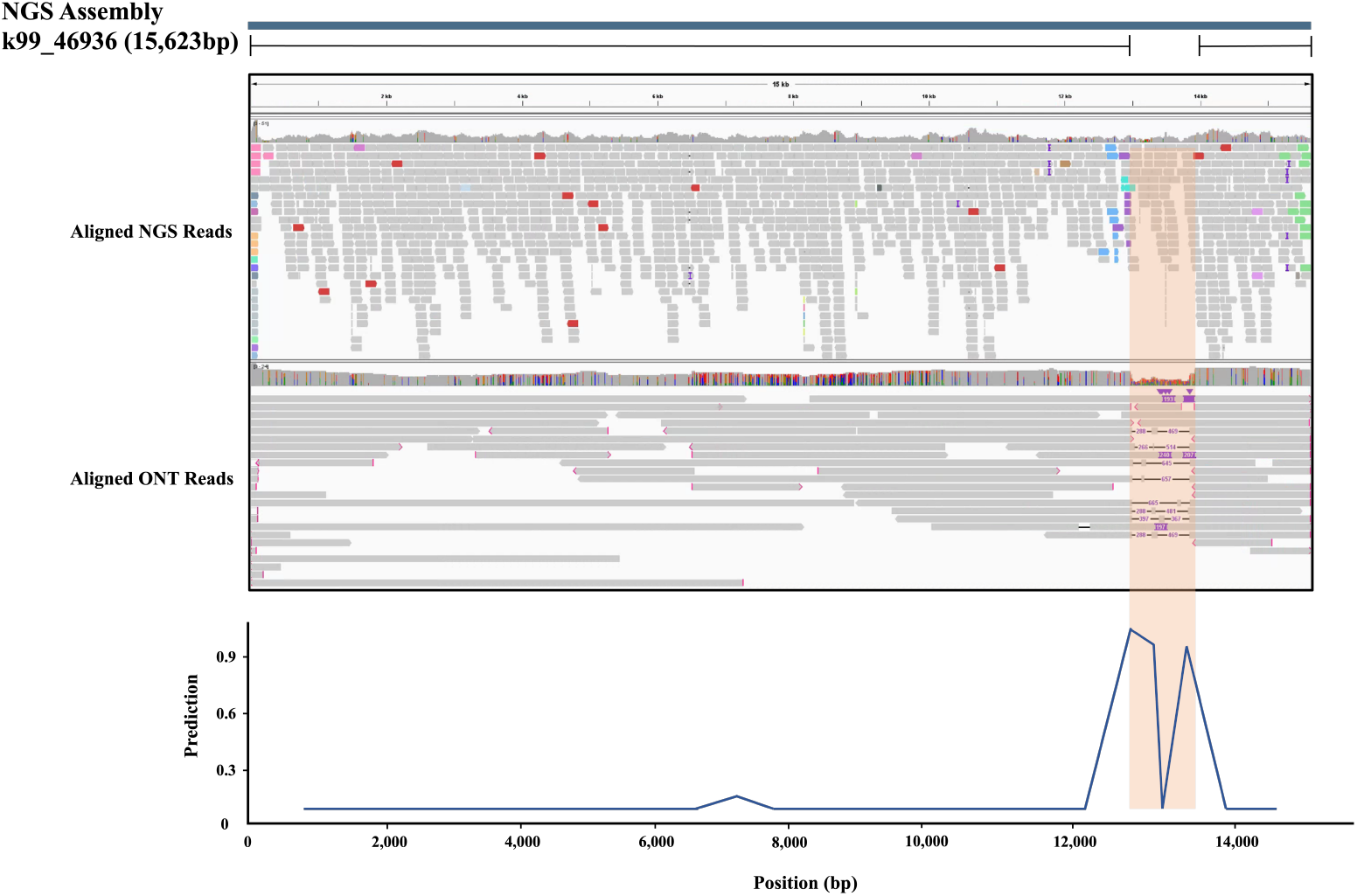
A case of long reads support misassembly in real metagenomic data

**Table 1:**
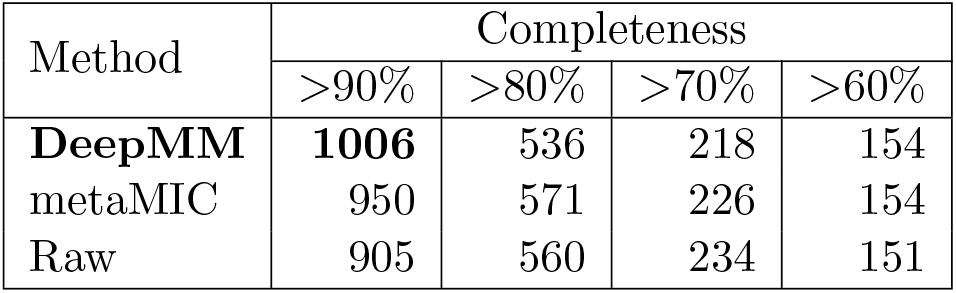
Increased the number of near-complete reconstructed bins in IBS-D data.

| Method | Completeness |  |  |  |
| --- | --- | --- | --- | --- |
|  | >90% | >80% | >70% | >60% |
| **DeepMM** | **1006** | 536 | 218 | 154 |
| metaMIC | 950 | 571 | 226 | 154 |
| Raw | 905 | 560 | 234 | 151 |

## 3 Methods

### 3.1 Alignment-purify processing

Current assemble algorithms using short reads may not resolve intra-species repetitive regions and inter-species conserved regions[17]. Aligning reads to assemblies with repetitive regions can introduce misleading alignment signals, potentially confounding existing evaluation methods. Consequently, these inaccuracies may lead certain correction techniques to erroneously correct assemblies by fragmenting them into shorter segments, thereby impairing the efficacy of binning tasks. Therefore, we design an alignment-purify processing to address this problem to avoid inaccuracies. Given a read aligned to a target assembly and its mate pair aligned to another assembly, we calculate the number of distinct assemblies to which the mate pair is aligned and exclude the target assembly, if this number exceeds two. Reads with low mapping quality(*<*40) and low alignment score(*<*100) are filtered out.

### 3.2 Multi-channel image generation

After alignment-purify processing, we constructed a sliding window from the beginning for each assembly, which skips the 10% base pair of each end due to bad assembly quality The skipping length will not be longer than 1k bp. We perform a *l* length sliding window to capture signals and define the following alignment signal functions to calculate each type of feature. We assign each position *i* of a window to a one-dimensional feature array *F* :

- The clipped reads feature

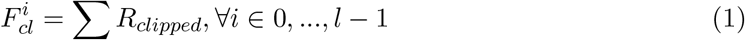

is counting the total number of clipped reads under position *i*.
- The inverted orientation reads feature

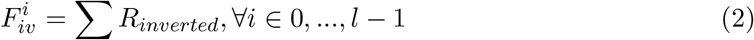

is counting the total number of read pairs that cover position *i* in the same orientation.
- The translocated read pairs feature

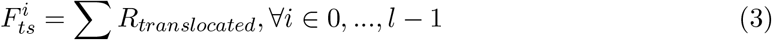

is counting the total number of reads that cover position *i* and its mate pair is aligned to different assemblies. These three features are normalized by read depth.
- Insert size feature

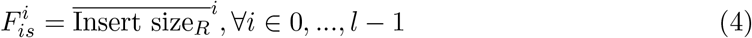

is represented by calculating the average insert size of read pairs that cover position *i* and is standardized to match different sequencer settings.
- The read depth feature

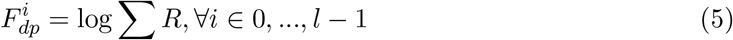

is counting the total number of reads aligned to position *i* and normalized by log function to align difference abundance.
- The read depth difference feature map

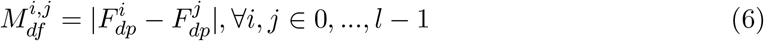

is computed with the difference of the number of reads that cover position *i* and *j* by the number of region reads.

Each array of features *F* will become a feature map *M* by assigning the values of *F* to the diagonal of *M*. Therefore, a 6 channel image is made up of 6 feature maps and passed to the neural network to predict the probability of the window if it contains a misassembly.

### 3.3 Misassembly window identifying and correction

The threshold of misassembly window prediction is 0.9. If an assembly contains multiple misassembly windows, we select the window with the highest prediction score. For this selected window, we identify the locations of clipped points from clipped reads and assign their aggregated counts to the corresponding positions on the assembly sequence. We then split the misassembly at the breakpoint with the highest number of clipped points. We only correct misassemblies in windows that include translocated read pairs and achieve the highest prediction score among all windows to prevent the generation of excessively fragmented sequences.

### 3.4 Dataset

The training datasets contain 100 metagenomic communities that were generated in the same way as metaMIC. In more detail, 1000 complete genome sequences from the Genome Taxonomy Database release 207[18] were used for simulating training datasets as metaMIC did. Then MGSIM[7] was used to create the 100 replicate metagenomic communities in which 10^7^ Illumina HiSeq2500 150bp paired-end reads were simulated per metagenome with the ART-defined default error distribution[19]. The abundance of each genome is sampled based on lognormal distributions, which are *Lognormal*(5, 2), (10, 1), (10, 2) and (10, 05), used in metaMIC and ResMiCo. Assemblies are constructed with MEGAHIT[10], and the length *<*3000 bp assemblies were removed. We split the dataset into two parts: 80% of the data was used to train the model and 20% of the data was used for model evaluation, where the evaluation set will not be seen or used during the training process. The label of each contig is provided by MetaQUAST[11] (parameters: –min-contig 1000 -extensive-mis-size 100). We selected the window that none of *F*_*bp*_, *F*_*iv*_, and *F*_*ts*_ is empty as a candidate window. For misassembled assemblies, we constructed the candidate window based directly on the breakpoint reported by MetaQUAST. The breakpoint was designated as the midpoint of the window. For correct assemblies, we conducted a sliding window approach to identify the candidate window. However, some assemblies with small gaps were also reported as correct by MetaQUAST, which could potentially mislead the model. Therefore, we only considered assemblies that exhibited 100% mapping to their reference genome sequences, as determined by the output files of MetaQUAST.

To evaluate the performance of different tools, five simulated datasets from CAMI and one real dataset are used to compare DeepMM and other competing tools in identifying misassemblies. All assemblies are mainly assembled by MEGAHIT if no notice. For fair evaluation, we balanced the number of misassemblies and correct assemblies in metaSPAdes assembled dataset as metaSPAdes produce less miassemblies. Only assemblies longer than 5000bp are considered as the majority of misassemblies that occur in assemblies are longer than 5000bp[5].

### 3.5 Data Augmentation

For each candidate window, we generated a total of five views: one original view and four augmented views. The augmented views were created by: (1) shifting the candidate window based on the midpoint to the left and right randomly, (2) sub-sampling the number of aligned reads with a ratio range from 0.6 to 1, and reconstructing the window using the reduced read sets, and (3) creating a reversed version of the original candidate window.

### 3.6 Contrastive learning

Contrastive learning derives instance representations through unsupervised proxy tasks by optimizing contrastive losses[20]. This process brings representations of similar instances closer together while pushing apart those of dissimilar instances. The determination of similarity is based on the specific unsupervised proxy task. In our identification task, different views from the same original window during the augmentation step are considered similar instances.

To effectively learn general visual misassembly representations, we applied SimCLR[21], an approach in contrastive learning. SimCLR learns representations by maximizing agreement between differently augmented views of the same data example using a contrastive loss in the latent space. The augmented images are encoded via an encoder network *f* (·) (a ResNet[22]) to generate representations. We utilized the normalized temperature-scaled cross-entropy (NT-Xent) loss function [23] as the objective function similar to COMEBin [24]. Each batch has *N* original windows, and each window has a total of *V* views. In each batch, the *V* (*V* = 5) views of each window are mutually positive samples, and all views from other windows are mutually negative samples. The embedding loss function for each batch is defined as follows. Let

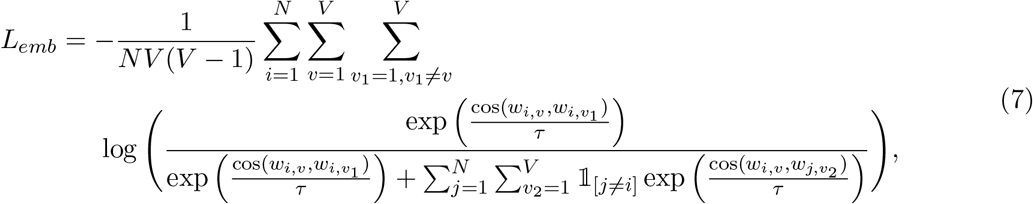

where *w*_*i,v*_ and 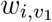 denotes the representation of the *v*-th and *v*_1_-th view of the *i*-th original window and *τ* is a hyper-parameter, which represents the temperature coefficient used to adjust the emphasis on similar negative samples. The indicator function𝟙_[*j*≠*i*]_ is defined to be one if and only if *j*≠ *i*.

The representations generated by *f* (*·*) are subsequently passed to an MLP projection head for misassembly window classification. A cross-entropy loss function is employed for this prediction task and the final loss function is

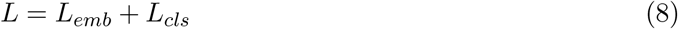

### 3.7 Model training and fine-tuning

We utilized ResNet-50 as the default model architecture due to its simplicity and superior feature learning capability. During training, we set the batch size to 64, the learning rate to 0.0003, the parameter *τ* to 0.07, and the output dimension of *f* (·) to 128. Model training was conducted using four Nvidia A100 GPUs, each with 80 GB of memory. We also trained DeepMAsED, metaMIC, and ResMiCo, and used the best performance between the re-trained and default models for fair evaluation.

Since reads may vary across different sequencers and inconsistent alignment features can affect model performance, we offer a fine-tuning process to enhance generalization capability. We use ART to generate read profiles and then employ MGSIM to simulate fine-tuning datasets. Insert size parameters can be found in sequencer documentation. For example, the CAMI dataset [25] has a distinct read profile compared to our simulation tool. Therefore, we generated a supplementary dataset simulated with the CAMI read profile characteristics for model fine-tuning. This dataset comprises 30 metagenomic communities, with the abundance of each genome sampled from *lognoram*(10, 1). All other parameters were consistent with those used for the pre-training dataset. During fine-tuning, we only adjusted the MLP projection head, using a batch size of 128 and a learning rate of 0.001 on a single Tesla V100 GPU with 40 GB of memory.

## 4 Discussion

We present DeepMM, a novel tool designed for the identification and correction of metagenomic misassemblies. By transforming various alignment features into a multi-channel image and leveraging contrastive learning to align different views of a misassembly, DeepMM achieves superior accuracy in misassembly identification. Higher AUPRC scores on simulated datasets and better improved binning results demonstrate the robust generalizability of DeepMM across different types of sequencing data. One of the key strengths of DeepMM is its fine-tuning process, which enables the model to adapt to data generated by various sequencers and assemblers, ensuring consistent performance across a wide range of datasets. In the future, we will consider employing DeepMM to other types of data, such as archaea, eukaryotes, or viruses. Misassembly correction process in DeepMM relies on splitting misassemblies into fragmented sequences. This poses a challenge for current binning methods, as fragmented sequences can complicate the reconstruction of MAGs[13]. Although our results show that DeepMM can enhance binning outcomes in near-complete MAGs, the quality of reconstructed MAGs with completeness *>*80% and *>*70% could be further improved by developing methods to connect broken sequences before the binning task.

## 5 Acknowledgements

The design of the study and the collection, analysis and interpretation of the data were partially supported by the Young Collaborative Research grant (no. C2004-23Y), HMRF (grant no. 11221026),

## 6 code availablity

The source code is freely available under the MIT license at GitHub: https://github.com/ericcombiolab/DeepMM

## Notes

### Competing Interest Statement

The authors have declared no competing interest.

### Summary of Updates

I have updated the GitHub code link to the preprint and corrected some typos.

## References

[1] Zha, Y., Chong, H., Yang, P. & Ning, K. Microbial dark matter: from discovery to applications. Genomics, Proteomics and Bioinformatics 20, 867–881 (2022).

[2] Behjati, S. & Tarpey, P. S. What is next generation sequencing? Archives of Disease in Childhood-Education and Practice 98, 236–238 (2013).

[3] Olson, N. D. et al. Metagenomic assembly through the lens of validation: recent advances in assessing and improving the quality of genomes assembled from metagenomes. Briefings in bioinformatics 20, 1140–1150 (2019).

[4] Slatko, B. E., Gardner, A. F. & Ausubel, F. M. Overview of next-generation sequencing technologies. Current protocols in molecular biology 122, e59 (2018).

[5] Lai, S. et al. metamic: reference-free misassembly identification and correction of de novo metagenomic assemblies. Genome Biology 23, 242 (2022).

[6] Clark, S. C., Egan, R., Frazier, P. I. & Wang, Z. Ale: a generic assembly likelihood evaluation framework for assessing the accuracy of genome and metagenome assemblies. Bioinformatics 29, 435–443 (2013).

[7] Mineeva, O., Rojas-Carulla, M., Ley, R. E., Schölkopf, B. & Youngblut, N. D. Deepmased: evaluating the quality of metagenomic assemblies. Bioinformatics 36, 3011–3017 (2020).

[8] Mineeva, O. et al. Resmico: Increasing the quality of metagenome-assembled genomes with deep learning. PLOS Computational Biology 19, e1011001 (2023).

[9] Sczyrba, A. et al. Critical assessment of metagenome interpretation—a benchmark of metagenomics software. Nature methods 14, 1063–1071 (2017).

[10] Li, D., Liu, C.-M., Luo, R., Sadakane, K. & Lam, T.-W. Megahit: an ultra-fast single-node solution for large and complex metagenomics assembly via succinct de bruijn graph. Bioinformatics 31, 1674–1676 (2015).

[11] Gurevich, A., Saveliev, V., Vyahhi, N. & Tesler, G. Quast: quality assessment tool for genome assemblies. Bioinformatics 29, 1072–1075 (2013).

[12] Nurk, S., Meleshko, D., Korobeynikov, A. & Pevzner, P. A. metaspades: a new versatile metagenomic assembler. Genome research 27, 824–834 (2017).

[13] Pan, S., Zhao, X.-M. & Coelho, L. P. Semibin2: self-supervised contrastive learning leads to better mags for short-and long-read sequencing. Bioinformatics 39, i21–i29 (2023).

[14] Chklovski, A., Parks, D. H., Woodcroft, B. J. & Tyson, G. W. Checkm2: a rapid, scalable and accurate tool for assessing microbial genome quality using machine learning. Nature Methods 20, 1203–1212 (2023).

[15] Zhao, L. et al. A clostridia-rich microbiota enhances bile acid excretion in diarrhea-predominant irritable bowel syndrome. The Journal of clinical investigation 130, 438–450 (2024).

[16] Chen, L. et al. Short-and long-read metagenomics expand individualized structural variations in gut microbiomes. Nature communications 13, 3175 (2022).

[17] Zhang, Z. et al. Exploring high-quality microbial genomes by assembling short-reads with long-range connectivity. Nature Communications 15, 4631 (2024).

[18] Parks, D. H. et al. Gtdb: an ongoing census of bacterial and archaeal diversity through a phylogenetically consistent, rank normalized and complete genome-based taxonomy. Nucleic acids research 50, D785–D794 (2022).

[19] Huang, W., Li, L., Myers, J. R. & Marth, G. T. Art: a next-generation sequencing read simulator. Bioinformatics 28, 593–594 (2012).

[20] He, K., Fan, H., Wu, Y., Xie, S. & Girshick, R. Momentum contrast for unsupervised visual representation learning. In Proceedings of the IEEE/CVF conference on computer vision and pattern recognition, 9729–9738 (2020).

[21] Chen, T., Kornblith, S., Norouzi, M. & Hinton, G. A simple framework for contrastive learning of visual representations. In International conference on machine learning, 1597–1607 (PMLR, 2020).

[22] He, K., Zhang, X., Ren, S. & Sun, J. Deep residual learning for image recognition. In Proceedings of the IEEE conference on computer vision and pattern recognition, 770–778 (2016).

[23] Caron, M. et al. Unsupervised learning of visual features by contrasting cluster assignments. Advances in neural information processing systems 33, 9912–9924 (2020).

[24] Wang, Z. et al. Effective binning of metagenomic contigs using contrastive multi-view representation learning. Nature Communications 15, 585 (2024).

[25] Marotz, C. et al. Evaluation of the effect of storage methods on fecal, saliva, and skin microbiome composition. msystems 6: e01329–20 (2021).

